# MaveRegistry: a collaboration platform for multiplexed assays of variant effect

**DOI:** 10.1101/2020.10.14.339499

**Authors:** Da Kuang, Jochen Weile, Nishka Kishore, Alan F. Rubin, Stanley Fields, Douglas M. Fowler, Frederick P. Roth

**Affiliations:** Donnelly Centre, University of Toronto, Toronto, M5S 3E1, Canada; Department of Molecular Genetics, University of Toronto, Toronto, M5S 1A8, Canada; Lunenfeld-Tanenbaum Research Institute, Sinai Health System, Toronto, M5G 1X5, Canada; Department of Computer Science, University of Toronto, Toronto, M5T 3A1, Canada; Bioinformatics Division, The Walter and Eliza Hall Institute of Medical Research, Parkville, VIC, 3052, Australia; Department of Medical Biology, University of Melbourne, Melbourne, VIC, 3010, Australia; Bioinformatics and Cancer Genomics Laboratory, Peter MacCallum Cancer Centre, Melbourne, VIC, 3000, Australia; Department of Medicine, University of Washington, Seattle, WA, 98195, USA; Department of Genome Sciences, University of Washington, Seattle, WA, 98195, USA; Department of Bioengineering, University of Washington, Seattle, WA, 98105, USA

## Abstract

**Summary:** Multiplexed assays of variant effect (MAVEs) are capable of experimentally testing all possible single nucleotide or amino acid variants in selected genomic regions, generating ‘variant effect maps’, which provide biochemical insight and functional evidence to enable more rapid and accurate clinical interpretation of human variation. Because the international community applying MAVE approaches is growing rapidly, we developed the online MaveRegistry platform to catalyze collaboration, reduce redundant efforts, allow stakeholders to nominate targets, and enable tracking and sharing of progress on ongoing MAVE projects.

**Availability and implementation:** https://registry.varianteffect.org

## Introduction

Germline genetic testing is increasingly carried out for patients with an elevated risk of hereditary diseases such as breast and ovarian cancer (Jones *et al.*, 2019). These tests frequently yield previously-unseen and often extremely rare genetic variants, amongst which it can be difficult to identify the subsets of pathogenic and benign variants (Blazer *et al.*, 2015). Indeed, within the broadly-used ClinVar repository of clinically-observed human variation (Landrum *et al.*, 2018), over 50% of missense variants are classified as ‘variants of uncertain significance’ (VUS) (Weile and Roth, 2018; Starita *et al.*, 2017).

After a variant is initially interpreted as VUS, functional testing experiments, e.g. complementation (Lee and Nurse, 1987; Osborn and Miller, 2007) or *in vitro* biochemical activity assays (Guidugli *et al.*, 2014; Millot *et al.*, 2012), can provide additional functional evidence that, under the American College of Medical Genetics and Genomics/Association for Molecular Pathology (ACMG/AMP) guidelines (Richards *et al.*, 2015; Brnich *et al.*, 2019), might re-classify the variant into more ‘actionable’ categories, such as ‘likely pathogenic’ or ‘likely benign’. However, results from such a ‘reactive’ functional testing approach can arrive weeks or years after initial discovery of a variant.

Multiplexed assays of variant effect (MAVEs) are experiments that offer more economical and consistently-measured tests of variant function than single-variant functional assays. MAVEs are generally conducted via a selection whose outcome depends on variant function. The selection is imposed on a heterogeneous culture of cells bearing different variants, and the frequency of these variants is tracked using next-generation sequencing (Starita *et al.*, 2017; Weile and Roth, 2018). MAVEs are inherently ‘proactive’: By testing the effect of nearly all possible single nucleotide or amino acid variants in a genomic region of interest, they provide functional evidence both for previously-observed variants and for the variants that will be observed in years to come.

The number of studies applying MAVE methods has grown in recent years (Weile and Roth, 2018; Esposito *et al.*, 2019), with nearly one hundred research labs involved thus far. This number is expected to grow, as national (Impact of Genomic Variation on Function (IGVF) Consortium) and international (Atlas of Variant Effects (AVE) Alliance, https://varianteffect.org) efforts are launched to systematically measure variant effects. To facilitate collaboration and communication at this scale, we created the MaveRegistry. This collaborative resource aims to serve multiple types of stakeholder, including interested researchers, clinicians, patients, patient advocates and funders. MaveRegistry enables users to: 1) browse MAVE studies posted to the MaveRegistry; 2) nominate targets for which a variant effect map would be beneficial; 3) report and follow progress on ongoing or published MAVE studies; and 4) follow progress updates from research teams. To facilitate sharing of progress of ongoing MAVE studies, we designed a role-based permission model, extending some functionality to anonymous users while making additional functionality available to registered members, project leads, and approved project and team followers.

### MaveRegistry promotes data sharing with a role-based permission model

The practice of sharing progress on unpublished projects online remains relatively unusual in academic research, but there are benefits and precedents to this practice. For example, a resource developed to share progress on mouse embryonic stem cells knocked out for particular target genes (Austin et al. 2004) has helped to inform the recipient community while avoiding duplicated efforts.

Sharing progress or data on unpublished projects is often a double-edged sword (Theologis and Davis, 2004). Although making unpublished information available can foster collaboration and reduce time and resources spent on redundant efforts (Fischer and Zigmond, 2010), researchers tend to be cautious because of a fear of losing credit for a finding to a competitor. To encourage researchers to share their planning of MAVE studies before any data are generated, MaveRegistry needs to address this dilemma. We therefore designed a role-based permission model (Figure 1A) to enable users providing project updates to control who can access their information. These roles are “public,” “member,” “follower” and “funder.”

**Figure 1.**
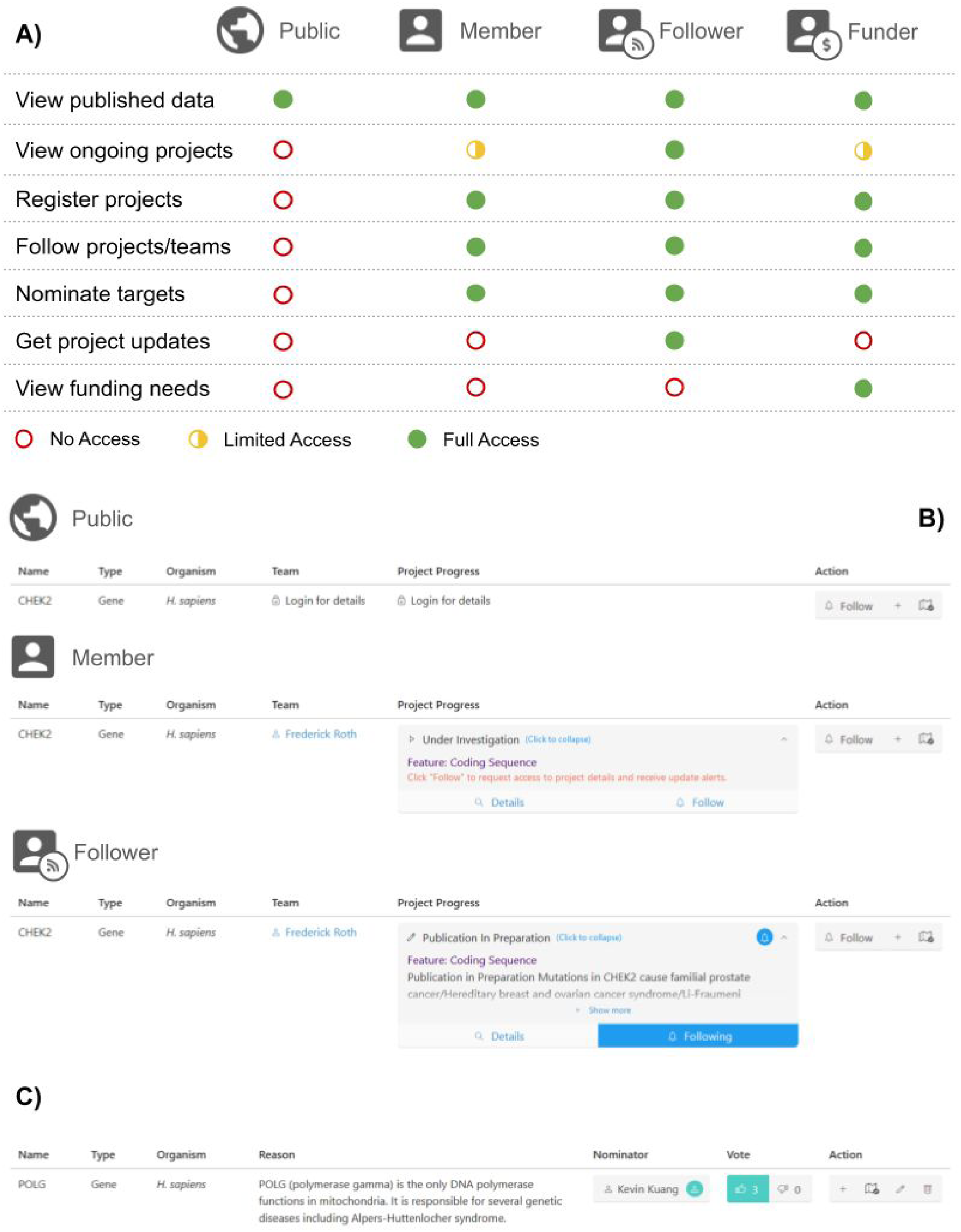
Core features of MaveRegistry. **A)** A role-based permission model enabling user-controllable access to information about ongoing variant effect mapping studies. **B)** Access to project information depends on the user’s role. **C)** A target nomination interface allows users to propose targets (i.e. protein-coding genes or other genomic regions) for variant effect mapping studies.

The general public (with a “public” role in Figure 1A), without logging in, can use MaveRegistry to access all information on MAVE projects that have been marked as ‘published’ by their depositors, which includes a brief summary of the project, with links for data retrieval (e.g. via MaveDB, an online repository of MAVE data (Esposito *et al.*, 2019)), and publications (e.g., via PubMed (NCBI Resource Coordinators, 2018)).

Once they have created an account and have thus become “members” (see Figure 1A), researchers engaged in MAVE studies can post their ongoing or published MAVE projects. Any member can request viewing and editing permission of posted MAVE projects. If approved by the depositor to ‘follow’ a project, members (now with a “follower” role in Figure 1A) can view project details and receive notifications when updates occur. Members may also request permission to follow research teams, so that they receive notifications when project status changes. Members affiliated with funding agencies may choose a “funder” role (Figure 1A), granting them access to the self-reported funding needs for each MAVE project, which depositors have the option to disclose. Figure 1B uses the *CHEK2* example to demonstrate which information is visible to different roles on MaveRegistry.

### Register and manage variant effect mapping projects

Members can register their ongoing and published variant effect mapping projects. If a genomic region (i.e. target element) has not been previously submitted, the submitting member will be prompted to enter basic information about the target, including its name, type (i.e. protein coding gene or other genomic region), and the relevant organism. Then, depositors provide contact information for at least one project lead, which will be made public to other members to facilitate direct peer-to-peer communication. At least one research team (which may include names of collaborators) is named for each project, enabling a “view projects by research team” function. To describe a project’s current status, depositors can currently select among six categories of project activity: “literature search”, “assay development/validation”, “MAVE data collection”, “MAVE data analysis”, “publication in preparation,” and “published”. For each activity, the start and end date (if the activity has been completed) can be entered. A brief free-text progress summary can also be provided, which is aided by a list of questions provided by MaveRegistry for consideration. To provide more flexibility to users, MaveRegistry enables depositors to link to external sources, e.g. relevant publications, code repositories, online lab notebooks or protocol resources.

When other members request to follow MAVE projects and teams, MaveRegistry notifies the corresponding depositors via on-site and email notifications. Through a moderation interface, depositors can review the reason-to-follow message submitted by follower, approve or reject each follow request, adjust follower’s permission (read-only or editable), or remove members from the list of followers. Depositors may also transfer ownership of submitted projects to other MaveRegistry members.

### Nominate targets for variant effect mapping

To foster proposal of and communication about new project ideas, we implemented a target nomination interface (Figure 1C) through which members can nominate targets for variant effect mapping, providing basic target information and their reasons for nomination. We expect that nominations might come from a wide range of stakeholders, ranging from patients, patient advocates and clinicians to researchers with an interest in the fundamentals of sequence-structure relationships. Other members can react to existing nominations by upvoting or downvoting. MAVE projects can be registered directly from the nomination interface, allowing the option to automatically fill in target information from this source. When the nominated target is a human protein-coding gene, users interested in pursuing a MAVE study may link to MaveQuest, an online resource for identifying potential functional assays and other information about clinical relevance (Kuang *et al.*, 2020).

We expect that the MaveRegistry platform will reduce unintentional competition, promote collaboration, and provide efficient communication about systematic variant effect mapping studies, while attracting additional interest from the general public and funding agencies.

## Acknowledgement

We appreciate help from members of the Roth lab, Lara Muffley and other members of the Center for the Multiplexed Assessment of Phenotype (a Center of Excellence in Genome Science of National Institutes of Health/National Human Genome Research Institute) and others who tested MaveRegistry and provided valuable feedback at various stages.

